# Different Contribution of the Monkey Prefrontal and Premotor Dorsal Cortex in Decision Making during a Transitive Inference task

**DOI:** 10.1101/2021.07.26.453792

**Authors:** S Ramawat, V Mione, F Di Bello, G Bardella, A Genovesio, P Pani, S Ferraina, E Brunamonti

**Affiliations:** Department of Physiology and Pharmacology, Sapienza University, Sapienza University of Rome, P.le Aldo Moro 5, 00185 Rome; PhD Program in Behavioral Neuroscience, Sapienza University of Rome, P.le Aldo Moro 5, 00185 Rome

**Keywords:** transitive inference task, prefrontal cortex, premotor cortex, monkey, decision making

## Abstract

Several studies have reported similar neural modulations between brain areas of the frontal cortex, such as the dorsolateral prefrontal (DLPFC) and the premotor dorsal (PMd) cortex, in tasks requiring encoding of the abstract rules for selecting the proper action. Here we compared the neuronal modulation of the DLPFC and PMd of monkeys trained to choose the higher rank from a pair of abstract images (target item), selected from an arbitrarily rank-ordered set (A>B>C>D>E>F) in the context of a transitive inference task. Once acquired by trial-and-error, the ordinal relationship between pairs of adjacent images (i.e., A>B; B>C; C>D; D>E; E>F), monkeys were tested in indicating the ordinal relation between items of the list not paired during learning. During these decisions, we observed that the choice accuracy increased and the reaction time decreased as the rank difference between the compared items enhanced. This result is in line with the hypothesis that after learning, the monkeys built an abstract mental representation of the ranked items, where rank comparisons correspond to the items’ position comparison on this representation. In both brain areas, we observed higher neuronal activity when the target item appeared in a specific location on the screen with respect to the opposite position and that this difference was particularly enhanced at lower degrees of difficulty. By comparing the time evolution of the activity of the two areas, we observed that the neural encoding of target item spatial position occurred earlier in the DLPFC than in the PMd.

## INTRODUCTION

When reaching a target, visual information about the target properties and location must be transformed into spatially oriented motor acts. Cortical circuits behind these transformations embed neurons that show spatial preference, i.e., higher neuronal activity for a specific target position compared to others (Wise et al., 1992; Caminiti et al., 1998; Ferraina et al., 2001; Lebedev and Wise, 2001). When more potential targets are simultaneously presented, neurons with different spatial preferences compete for signaling the proper target location and reaching direction (Cisek and Kalaska, 2005). An open question is whether and how perceptual or cognitive factors affect this decision-making process and subtending neuronal activity. The manipulation of perceptual variables, such as visual discriminability, demonstrates that the difficulty in encoding the target position was reflected in less sharp spatial preference in different brain areas (Coallier and Kalaska, 2014; Coallier et al., 2015; Chandrasekaran et al., 2017). However, how cognitive variables influence spatial preference-related activity in brain areas involved in visuomotor transformation is still poorly investigated. It has been recently demonstrated that in the dorsal premotor cortex (PMd), a brain area with a key role in visuomotor transformation in primates (Johnson et al., 1996; Boussaoud et al., 1998), neurons express their spatial preference depending on the cognitive difficulty in selecting the target when simultaneously presented with a non-target (Mione et al., 2020). In the quoted work, by using a Transitive Inference (TI) protocol, monkeys were first required to create a mental representation of a rank-ordered set of visual items as A> B> C> D> E > F and then to use this representation to select the higher-ranking item (target item) within all possible pairs by performing a reaching movement. In different trials, the target item was randomly presented on the left or the right side of a computer screen and paired with a non-target-item presented at the opposite spatial position. Within this experimental design, cognitive difficulty was modulated by pairing target and non-target items as a function of their proximity to the item in the mental representation of their ordinal position. The degree of proximity of pairs of items is quantified by their difference in rank and referred to as the symbolic distance (SDist) effect at the behavioral level. Depending on the symbolic distance, target selectivity for item B - for example - was easier when paired with non-target item E (higher distance between items on the mental representation corresponding to SDist = 3) and more difficult when paired with non-target item C (lower distance between items on the mental representation corresponding to SDist = 1). The authors found that this cognitive difficulty affected the neural spatial preference (spatial selectivity) in PMd.

Several neurophysiology studies have revealed comparable neural activation between the PMd and DLPFC while encoding a given task-relevant variable, thus suggesting an overlapping competence of the two brain areas (Cromer et al., 2011; Yamagata et al., 2012; Fine and Hayden, 2021). It is known that the prefrontal cortex can flexibly encode different task-relevant variables (Donahue and Lee, 2015; Fusi et al., 2016; Astrand et al., 2020), including the position of the target in oculomotor delayed visual search and delayed-response tasks (Iba and Sawaguchi, 2002). Here, we first asked if target position selectivity of the DLPFC emerges during the decision process of a TI task, and then we tested for differences between the DLPFC and PMd.

We observed that the spatial selectivity of both DLPFC and PMd neurons was modulated by cognitive difficulty in target selectivity. Importantly, the comparison between the time evolution of neuronal activity in the two brain areas revealed that the selection of target position occurred with different timings in the DLPFC and PMd. In this work, in line with previous studies (Cromer et al., 2011; Yamagata et al., 2012), we support the idea of a tight interplay between the lateral PFC and PMd in the target position encoding for action execution.

## MATERIALS AND METHODS

### Subjects and Data Acquisition

#### Animals

Neuronal activity was recorded from the DLPFC and the PMd of three male rhesus monkeys (Macaca mulatta), weighing 5.50 kg (Monkey 1), 6.50 kg (Monkey 2), and 10.0 kg (Monkey 3) respectively, while performing a TI task. Monkey 1 and Monkey 2 were used to record the data from DLPFC as reported in (Brunamonti et al., 2016); Monkey 1 and a third monkey, Monkey 3 (Mione et al., 2020; Monkey 3 is referred as Monkey 2 in the referred article) were used for the PMd recording.

#### DLPFC Recording

In Monkey 1 and Monkey 2, the cells in the dorsal area of the prefrontal cortex (DLPFC; Figure S1) were targeted, and the neural activity from these cells was recorded extracellularly using a five-channel multielectrode recording system (Thomas Recording, Germany) acutely and in different sessions. The recording chambers were surgically implanted in the left frontal lobe at stereotaxic coordinates: anterior-32 and lateral-19 in Monkey 1 and anterior-30 and lateral-18 in Monkey 2 along with head restraining devices (Brunamonti et al., 2016). At the end of the neurophysiological experiment, the location of the electrodes was confirmed by visual examination following the surgical opening of dura for the implantation of a chronic array in Monkey 1 (given the following recording from PMd neurons), while in Monkey 2, the electrodes’ location was confirmed using a structural MRI scan.

#### PMd Recording

Neural activity was recorded extracellularly from the left PMd of Monkey 1 and Monkey 3 (Mione et al., 2020; Figure S1) using chronic electrode arrays (96-channels; Blackrock Microsystems, Salt Lake City, Utah).

The surgical procedures in all three monkeys were performed under aseptic techniques while keeping the animal under general anesthesia (1-3% isloflurane-oxygen, to effect).

The monkeys were housed and cared for following the European (Directive 2010/63/EU) and Italian (D.L. 26/2014) laws, regulating the use of nonhuman primates in scientific research. The Italian Ministry of Health approved the research protocol. Housing conditions and experimental procedures were in line with the Weatherall report (use of nonhuman primates in research).

#### Behavioral data recording and task implementation

The task was controlled using the Cortex Software package (https://nimh.nih.gov/) and administered by presenting the task stimuli on a touchscreen (MicroTouch, sampling rate of 200 Hz) connected to the computer through a serial port to detect the response. An RX6 TDT recording system (Tucker-Davis Technologies, Alachua, FL, USA) was employed during the DLPFC recording sessions, which was synchronized to the behavioral events to detect and record neuronal activity during each trial. During the PMd recording sessions, an RZ2 TDT system (Tucker-Davis Technologies, Alachua, FL, USA) was used, since a higher number of simultaneously recorded channels were dealt with.

### Test stimuli and task design

For this study, at the beginning of every session, six stimulus images were randomly selected from a database of 80 abstract black and white images (16° X 16° visual angle, bitmaps) and arbitrarily ordered to form a ranked series (Figure 1A). To avert the familiarity of the stimulus/rank association, any of the stimuli were not repeated or assigned to the same rank for a significant number of consequent sessions (Brunamonti et al., 2014, 2016; Mione et al., 2020). The monkeys were trained to learn the relationship among all the items of this ranked series.

**Figure 1.**
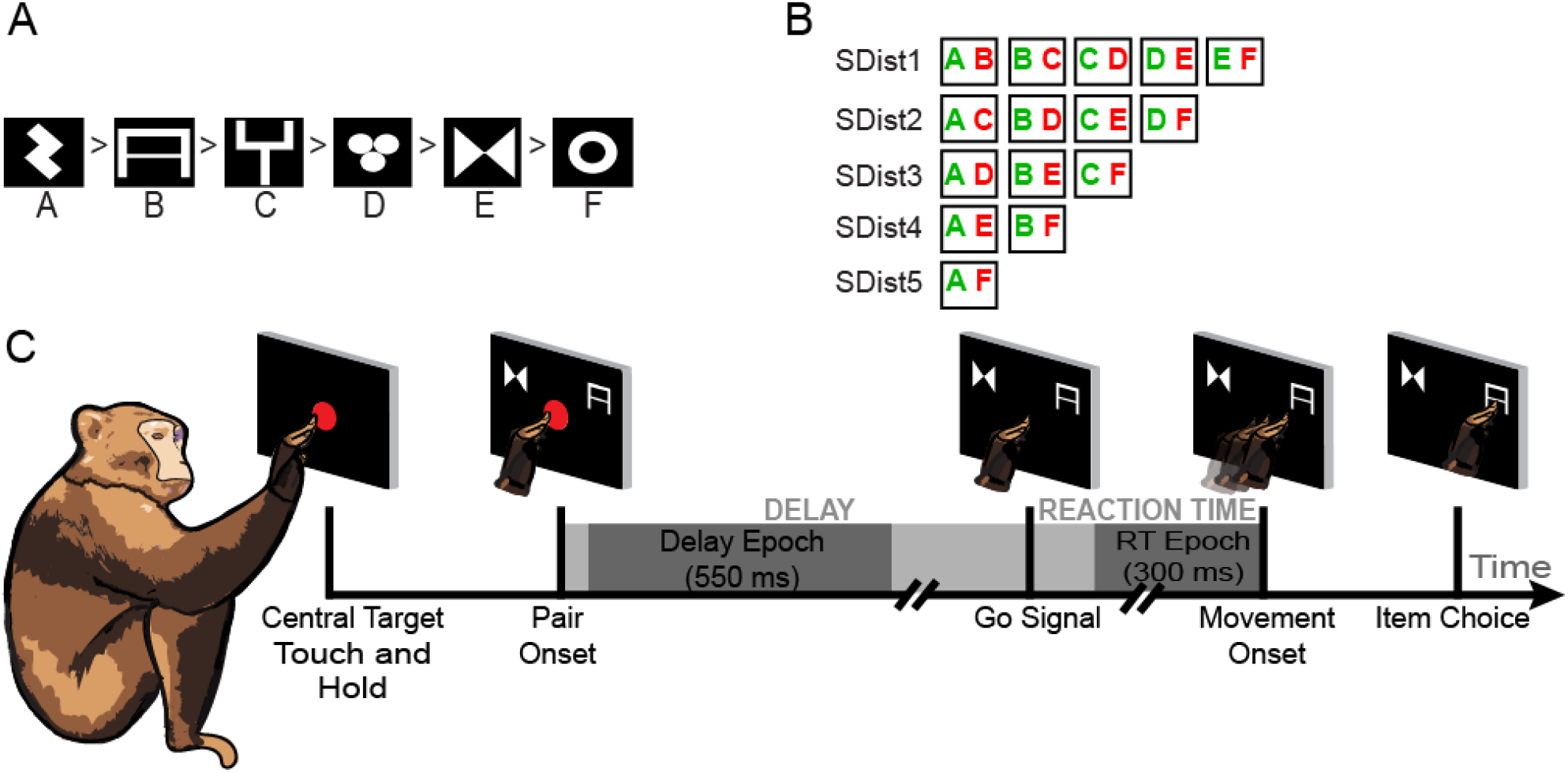
Task design and Analysis Epochs. **A.** An example of a 6-item ranked series chosen at the beginning of the experiment. The letters under each symbol are used here for illustrative purposes only. **B.** All the possible combinations of pairs from the items of the ranked series and their symbolic distances (SDist) associated with each combination; target items represented in green and non-target in red, following the link of each symbol with a letter as in A. **C.** Time course of a single trial during each session. The epochs used for analysis of neuronal activity are highlighted (dark grey).

Each experimental session (for both DLPFC and PMd neural recordings) comprised a learning phase, requiring the monkeys to learn the novel relationship between all the adjacent items (Figure 1B: SDist1) in the ranked series, and a test phase in which the monkeys were asked to infer the relationship between novel pairings of nonadjacent items (Figure 1B; SDists >1). The learning phase was accomplished by means of two different methods: 1) sequential learning and 2) a chained learning procedure, as reported in our previous works (Brunamonti et al., 2016; Mione et al., 2020). More specifically, in the sequential learning sessions, monkeys were required to progressively learn the reciprocal rank order of pairs of items adjacent in the sequence, while in the chained learning procedure, monkeys were required to link two lists of 3 rank-ordered items previously acquired by sequential learning.

Sequential learning was implemented in two steps: learning phase 1 and learning phase 2. During learning phase 1, the monkeys were presented with the pairs comprising adjacent items sequentially, and they were required to identify the higher ranking item by trial and error in blocks of 15 (Monkey 1: DLPFC recording) or 20 (Monkey 2: DLPFC recording; and Monkey 1 and Monkey 3 PMd recording) trials. Each block was repeated until the monkey achieved a performance of at least 90% (DLPFC recording) or 80% (PMd recording) for the pair. Once the desired performance was reached for every pair in the series, learning phase 2 followed, where the previously experienced pairs were presented in a random order in larger blocks of trials, and a different criterion (> 60% correct trials) was used.

In contrast, during the chained learning procedure, the six-item list was divided into two smaller three-item lists (A>B>C and D>E>F). The rank order of the two three-item lists was learned independently by the sequential learning method in separate blocks until the monkeys achieved a criterion of at least 80% correct trials. The two lists were then linked by learning the association C>D in blocks of 20 trials. The chained learning procedure allowed us to study the manipulation of the ranked order of the series according to the newly acquired information of the linking pair C>D; hence, the rank of the items in the second series was redefined in the unified list, supporting the behavioral strategy of inferential reasoning.

Both learning procedures allowed the monkeys to perform the test phase with a performance significantly above the chance level. We have previously reported that the performance in the test phase after the two different learning methods was comparable (Mione et al., 2020).

For the DLPFC experiment, all the sessions were recorded with Monkey 1 and Monkey 2 using sequential learning only (see Brunamonti et al., 2016 for further details), whereas during the PMd recording, Monkey 1 and Monkey 3 were tested for 7 sessions each using different learning procedures: chained-list learning in 6 (Monkey 1) and 4 (Monkey 3) sessions and sequential learning in 1 (Monkey 1) and 3 (Monkey 3) sessions (see Brunamonti et al., 2016; Mione et al., 2020 for further details).

In the test phase, all the possible pairs of items were presented in a random order (Figure 1B; SDist1 – SDist5). Each pair was presented at least 14 or 18 times during the DLPFC and PMd experiments, respectively, with an equal probability of the target item being presented on the left or the right position of the screen.

The time course of each trial was identical for the learning phase and the test phase during all the recording sessions from the DLPFC and PMd (Figure 1C). At the beginning of each trial, a red dot (central target) was presented at the center of the screen, and the monkey had to respond within 5 s by pressing a button fixed on the monkey chair (DLPFC sessions) or by touching the red dot on the screen (PMd sessions; as depicted in Figure 1C) until the appearance of a pair of items on the screen. The monkey was required to maintain the touch (or the button pressed) for a random variable Delay period (600-1200 ms) after the Pair Onset until the red spot disappeared from the screen (Go Signal). The Go Signal instructed the monkeys to select (to reach) the higher-ranking item on the screen to obtain the reward.

### Data Analysis

#### Selection of DLPFC and PMd recording sessions

One of the goals of the present work is to compare the temporal evolution of the neuronal encoding of the target position between the DLPFC and PMd while performing a TI task. Since this comparison was performed by analyzing the neuronal activity recorded in different experimental sessions, we first assessed that differences in the time evolution of the brain activity between the two areas were independent of different levels of accuracy and response time of monkeys in different sessions. To this aim, we evaluated the mean performance and the RTs of the DLPFC and PMd sessions from all three animals and observed that sessions from both experiments were characterized by significantly different RTs (Table 1). Additionally, we observed that the RTs of monkey 1 from the DLPFC (296 ms) and the PMd (393 ms) sessions were significantly lower than those of monkey 2 in the DLPFC sessions (386 ms) and monkey 3 in the PMd sessions (437 ms). To account for these differences, in the following analyses, data obtained from Monkey 1 were analyzed independently from those obtained in recording sessions with Monkey 2 and Monkey 3. To compensate for the differences between RTs from DLPFC and PMd sessions in each comparison, we calculated a range of RTs from PMd sessions (μ ± 3σ, where μ is the mean and σ is the standard deviation). As there were only 7 sessions recorded from the PMd of Monkey 1 and Monkey 3, we selected all the sessions recorded from PMd and selected a subsample of DLPFC sessions (14 sessions from Monkey 1 and 22 sessions from Monkey 2) conforming to the defined range of RTs. As a result of this selection, we obtained comparable RTs between Monkey 1 DLPFC and PMd sessions (all ps>0.05), Monkey 2 DLPFC, and Monkey 3 PMd sessions (all ps> 0.05; see Table 1 for details).

**Table 1:**
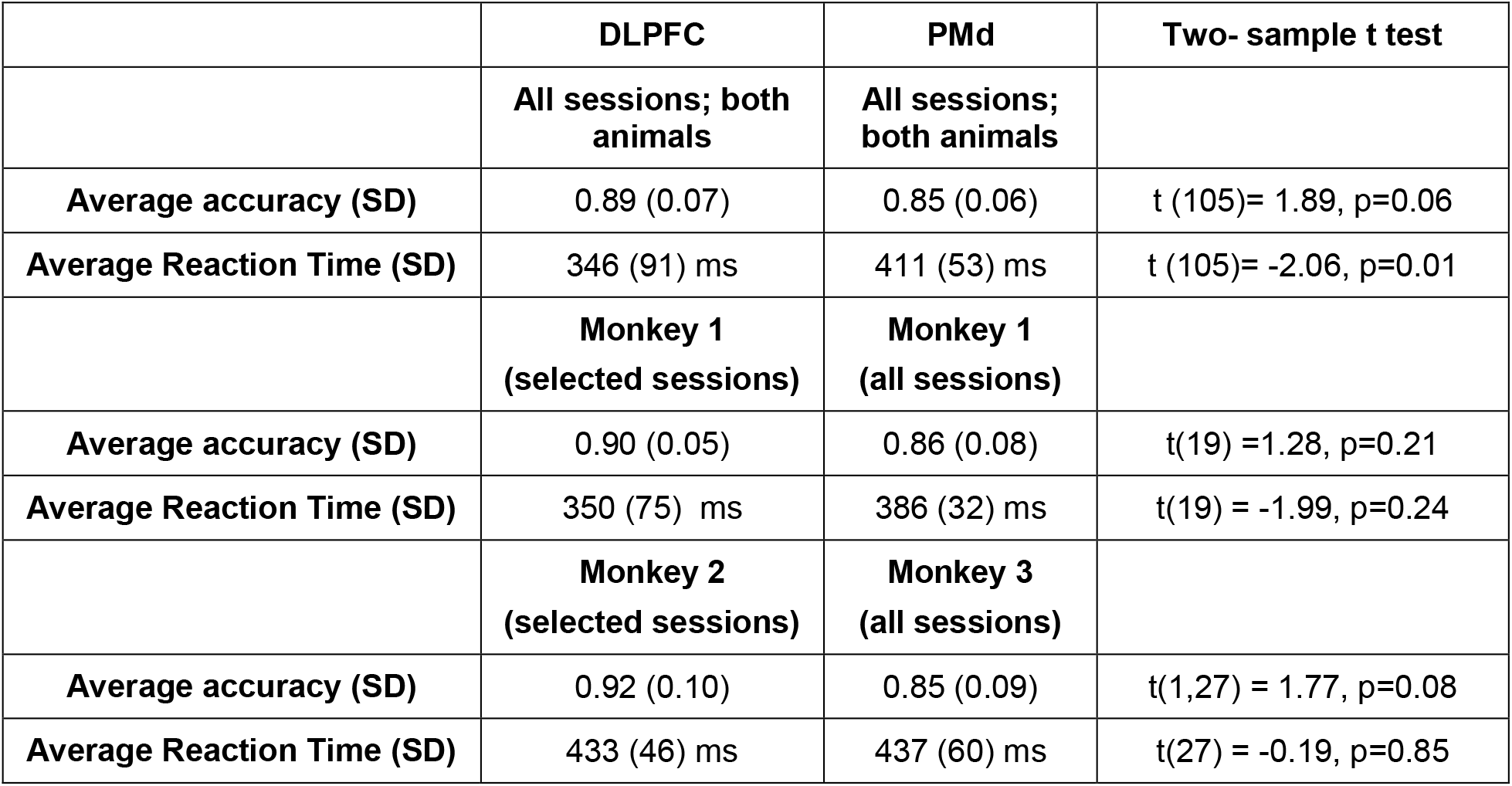
Comparative analysis of average behavior average behavior across PFC and PMd sessions, before and after selection of sessions; SD-Standard Deviation.

#### Behavioral correlates of TI at test

We first investigated whether the behavior during the selected sessions from DLPFC (Monkey 1: 14; Monkey 2: 22) and PMd (Monkey 1: 7; Monkey 2: 7) recordings was modulated by the SDist. This analysis was performed to assess whether the monkeys’ ability to select the target item over the simultaneously presented non-target item depends upon the distance of the two items in their mental representation. To this aim, we tested whether the probability of selecting the correct item significantly increased and the corresponding RT significantly decreased with increasing symbolic distance. To further investigate if the monkeys employed the mental models to solve the task during the selected sessions, we calculated the performance and the normalized reaction time for the selected DLPFC (Monkey 1: 14 and Monkey 2: 22) and all the PMd (Monkey 1: 7 and Monkey 3: 7) sessions for each pair comparison. Using a linear regression, we analysed if the performance and the RTs for individual item comparisons correlated with the SDist (Figure S2).

#### Neuronal correlates of TI in the test

We studied 141 neurons from the DLPFC (Monkey 1: 56, Monkey 2: 85) and 186 neurons (Monkey 1: 59, Monkey 3: 127) from the PMd obtained during the test phase of the TI task of the selected recording sessions. The data analysis was performed on trials in which the monkey correctly selected the position of the target item. Error trials in which the monkey selected the non-target items were not considered in the following analyses. To address whether the activity of DLPFC neurons correlated with the difficulty in selecting the position of the target item, we first studied spatial preference for the target that emerged in the neuronal activity of DLPFC, and then we tested whether this preference was modulated by the SDist. As a further step of the analysis, we compared the pattern of neuronal activity in the DLPFC and PMd to evaluate how it differed between the two brain areas during different epochs of the task. In doing so, we studied the neuronal activity from each neuron during two task epochs, 1) the Delay epoch, lasting from 50 ms to 600 ms after the Pair Onset; 2) the Reaction Time epoch, lasting for 300 ms before the Movement Onset (Figure 1C). For each neuron, we calculated the mean spike rate from the correct trials corresponding to each of the two target positions (left and right) at every SDist (SDist1-SDist5). Furthermore, we normalized the mean spike rate across different task conditions by applying a z score transformation.

To quantify the degree of preference of each neuron for the left or right target position, we first represented the neuronal activity as a point in a 2-dimensional space having the neuronal activity for the right and left positions of the target as x and y coordinates, respectively (Figure 3,4). Then, we computed the shortest distance (D_n_) of each point from the equality line (representing a preference or a lack of preference for any of the two target positions) as follows:

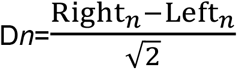

where Left_n_ and Right_n_ are the coordinates of the n^th^ neuron in the defined space, representing the normalized mean neuronal activity during the trials with the target located in the left and right positions, respectively. D_n_ represents the difference between the neuronal activity between these two target positions in this graphical context (Figure 3B), where the negative values represent the neuronal preference for the left target position and positive values for the right target position. A high magnitude of D_*n*_ (positive or negative) indicates a greater degree of preference for one of the two positions of the target, which we further utilized to identify the neurons showing a significant target position preference. Across the population of neurons from each area, we calculated the maximum range of variation during the two analysis epochs: D_population_ = (D_n_max_ – D_n_min_). Then, we defined a neuron to be selective for the left or the right target position if the corresponding positive or negative value D_n_ was lower (D_n_ < 0) or higher (D_n_ > 0) than 10% of the total range of excursion (D_population_). To study how the preference for the target position was modulated by the task difficulty, we first identified the DLPFC and PMd neurons displaying a target position selectivity at SDist5 (the easier condition of the task), and then we observed how this selectivity changed across other SDists. We used a linear model fit to test whether the target position selectivity was significantly modulated by the rising SDist for each target position (left and right).

**Figure 2.**
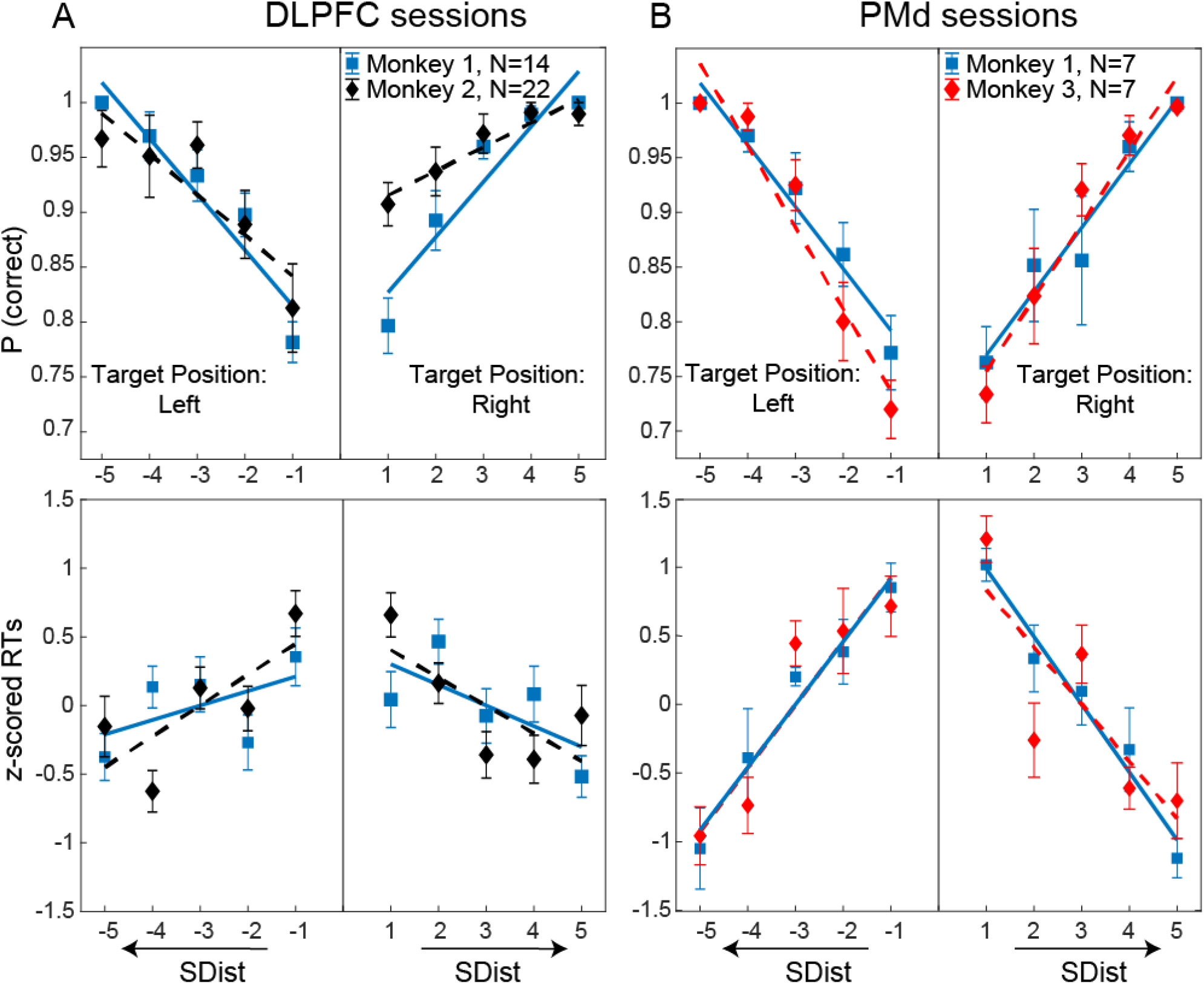
Influence of SDist on behavioral performance. Proportion of correct choices (top plots) and corresponding z scored RT (bottom plots) for target selection at every SDist when the target was presented on the left and the right position of the screen during ‘N’ recording sessions from DLPFC (A: Monkey 1 and Monkey 2) and PMd (B: Monkey 1 and Monkey 3). Squares and diamonds correspond to the average performance and z scored RT, respectively; across the sessions, vertical bars represent the S.E.M. Linear regression fit across sessions is reported in Table 2.

**Figure 3.**
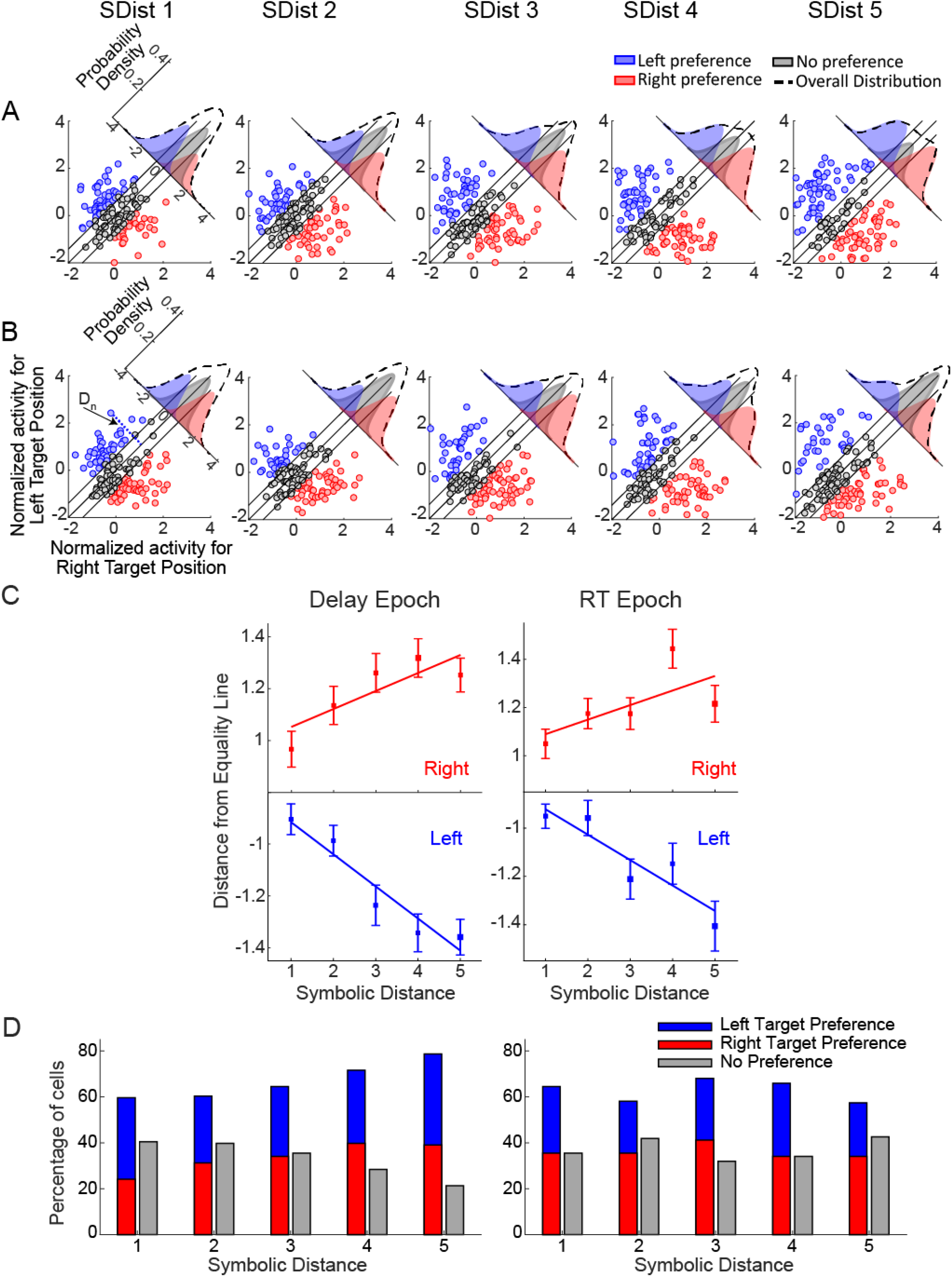
Encoding of target position in the population of DLPFC neurons. **A-B**, Scatter plots comparing the normalized mean neuronal activity from 141 DLPFC neurons during the delay (A) and RT epochs (B), calculated across all the SDists when the target was presented in the left (y-axis) and right (x-axis) positions on the screen. Each dot in the plots represents a single neuron. The preference for the left (blue dots) or right (red dots) target position of each neuron was detected as its distance (D_n_) from the equality line (diagonal in each plot). Neurons with a distance lower than 10% of the total range from the equality line were labeled nonselective for the target position (gray dots). The density plots (dotted line) on the top of each diagonal show the distribution of the spatial selectivity of neurons at each SDist, while the colored density plots represent the corresponding sub-distributions for the neurons showing selectivity for left, right, or no target preference. **C.** Mean distances of all the target position selective DLPFC neurons from the diagonal at each SDist for their preferred target positions (left or right), and their linear correlation with the SDist during the delay and RT epochs. They represent the strength of selectivity for left and right target positions exhibited collectively by the neurons, and the regression lines indicate a modulation of target position selectivity during delay (left: p<0.01 and right: p<0.01) and the RT epochs (left: p<0.01 and right: p<0.01). Vertical bars indicate the Standard Error of Mean. D. Percentage of neurons showing directional target preferences at varying choice difficulties.

**Figure 4:**
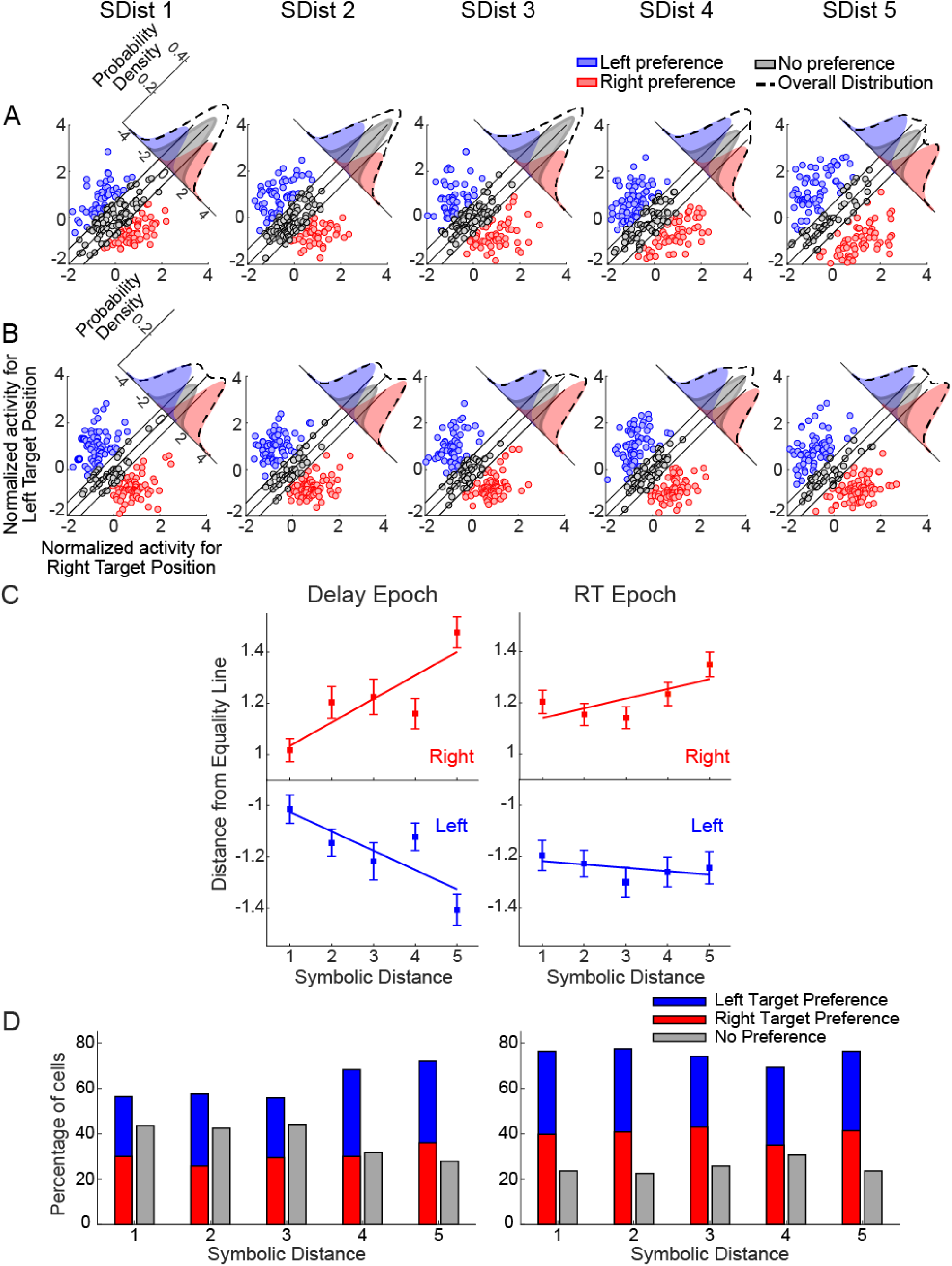
Encoding of target position in the population of PMd neurons. **A-B**, Scatter plots comparing the normalized mean neuronal activity for two target positions in 186 PMd neurons at each SDist during the Delay(A) and RT(B) epochs. The modulation of the neuronal activity is represented with the same codes as in Figure 3. **C.** The plots show the average strength of target position selectivity by overall target position selective PMd neurons at each SDist during the Delay and RT epochs for left and right target positions. It indicates a linear correlation between the strength of selectivity with the SDist during the Delay (left: p<0.01 and right: p<0.01), and a weaker modulation for the right target position and no modulation (p>0.05) for the left target position during RT by the SDist. **D.** Percentage of the PMd population showing target preferences at different SDists.

We applied a Receiver Operating Characteristic (ROC) analysis on neuronal activity recorded during the selected sessions to study the time course of target position selectivity in DLPFC and PMd (Thompson et al., 1996; Mione et al., 2020). Once the preferred target position of each neuron at SDist5 was identified, we computed the area under the ROC curve (auROC) in consecutive time bins in a time epoch starting 500 ms before to 1300 ms after the pair onset using a moving time window of 200 ms with a 20 ms step. Thus, for each time window of every neuron, we obtained a probabilistic measure of encoding the target when it was presented in the preferred position compared to the opposite position. We estimated the latency of target position discrimination by calculating the time when the auROC value crossed the threshold of 0.6 for 60 ms (3 consecutive time bins) in the trial time ranging from 80 ms to 1300 ms after the Pair Onset. We repeated this estimate for the comparisons at SDist lower than 5 to test if the latency of spatial discrimination changed in pairs comparisons of increasing degrees of difficulty in both, DLPFC and PMd. The auROC values were calculated for a balanced number of trials for all the task conditions, corresponding to the minimum number of correct trials across all the task conditions. On average, the number of trials for each position of the target was 17 (SD=5) for DLPFC sessions and 31 (SD=16) for PMd sessions.

To account for the differences in RT detected in Monkey 1 and the other two monkeys (see previous section), we performed latency estimates of target position selectivity in different groups of neurons: 1) neurons obtained by Monkey 1 in which the RT in DLPFC (47 neurons) and PMd (55 neurons) recording sessions was not significantly different; 2) neurons obtained by Monkey 2 in DLPFC (79 neurons) sessions and neurons recorded in Monkey 3 PMd (119 neurons) sessions with RT were not significantly different.

The latency of target position discrimination in neurons measured in Monkey 1 and those measured in Monkey 2 and Monkey 3 were analyzed separately. To test for the differences between latencies across different task conditions and different areas, we calculated the probability of observing latencies in the initial 75% of the distributions (< 900 ms). Consequently, we applied a Kolmogorov Smirnoff test to identify the inter-area differences between these probabilities if any. At test was used to compare the inter-area differences between the average latencies across the different SDists. To explore further how the neuronal populations of two areas encode target positions with varying difficulties, we computed the fraction of cells showing sustained coding for the target position (auROC>0.6 for more than 70% of the total time) as a function of SDist. The differences in the measures between the two areas were tested using a Kolmogorov Smirnoff test. All data analyses were performed using custom-made functions developed in MATLAB (The MathWorks) and Wolfram Mathematica.

## RESULTS

### Choice difficulty modulates the ability to select the target position during the TI task

The accuracy and RT for target position selection exhibited a significant SDist effect during the pair comparisons of the test phase in both the DLPFC and PMd recording sessions (Figure 2, Figure S2). A linear regression analysis revealed a significant increase in the proportion of correct to incorrect choices and a decrease in the RT with the rising SDist (see table 2 for details on the regression analysis) for each monkey. These results are in line with the hypothesis that, at the end of the learning phase, ranked items are arranged on a mental line, and the difficulty in pair comparisons at test depends on the relative proximity of the items’ representation on this line (Brunamonti et al., 2011, 2016; Mione et al., 2020).

**Table 2.**
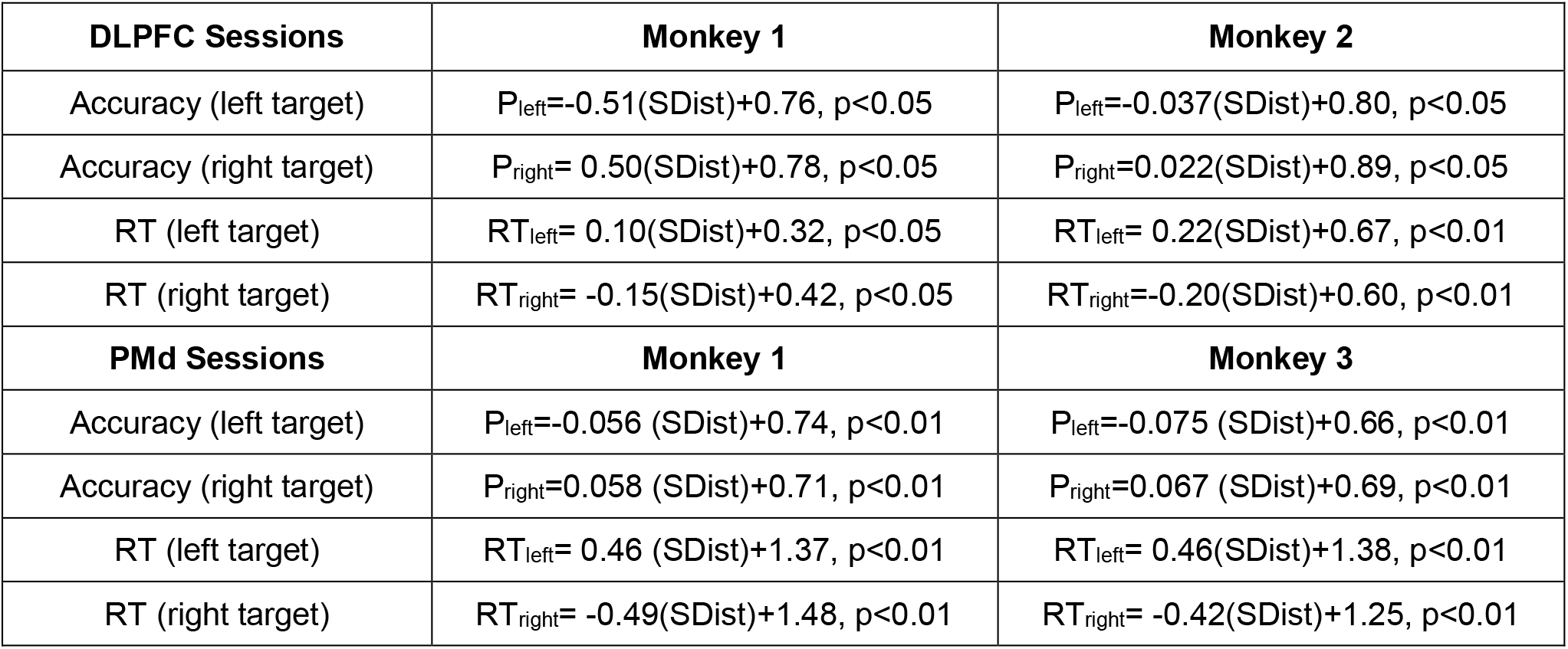
Linear Correlation Coefficients of SDist with behavior of each monkey during DLPFC and PMd recording sessions.

### Populations of DLPFC and PMd neurons encode the task difficulty while differentiating between the target positions

Here, we investigated whether the neuronal correlates of target position selectivity in the population of DLPFC (n=141) and PMd (n=186) neurons were modulated by the difficulty of the task.

Figures 3 and 4 display the scatterplots of the normalized neuronal activity in response to the presentation of the target on the left (y-axis) or right (x-axis) position on the display for each SDist during the Delay (Figures 3A and 4A) and Reaction Time (Figures 3B and 4B) analysis epochs. By representing the activity of each neuron as a point in such 2-dimensional space, we quantified the degree of preference of the target position as the distance from the equality line. According to this criterion, a neuron was classified as spatially selective if its distance from the equality line exceeded 10% of the total range of excursion of the distances calculated across all the neurons and SDists. DLPFC neurons were considered target selective for the left or right position of the target during the Delay epoch, if their distance from the equality line was lower than −0.49 or higher than 0.49, respectively. The same criterion in the RT epoch identified −0.51 and 0.51 as the threshold for left and right target selectivity. Similarly, a PMd neuron was distinguished to be spatially selective if the magnitude of the distance from the diagonal exceeded ±0.54 and ±0.46 during the Delay and RT epochs, respectively. For each neuron, we first identified the left (blue dots) or right (red dots) preference for the target position, and then we tested whether the encoding of the target location changed as the difficulty in pair comparisons gradually increased, i.e., during the pairs comparisons from SDist5 to SDist1.

We quantified the proportion of neurons exhibiting target position selectivity at each SDist. The colored density distributions on the top of diagonals in both Figures 3 and 4 show that the proportion of target position-selective neurons identified in the DLPFC and PMd during the Delay epoch gradually decreased as the difficulty in pair comparisons increased (see Figure 3D and 4D for details). At the same time, the proportion of neurons not showing a significant target position selectivity gradually increased from SDist5 to SDist1 (gray density distributions). This effect was not observed during the RT epoch in either area.

We further investigated the strength of target position selectivity as a function of the degree of difficulty in comparing the pairs of items by quantifying the average distance from the equality line for each SDist (leftward and rightward shifting of the blue and red sub-distributions in Figure 3 A-B and Figure 4 A-B). By means of regression analysis, we detected that for both left and right target position selective neurons, the distance from the equality line significantly increased with the increasing SDist in DLPFC neurons during the delay and RT epochs for each of the target positions (left and right: p<0.01; Figure 3C). However, in the PMd neurons (Figure 4C), this effect was mainly detected only in the delay epoch (left and right: p<0.01) but not in the RT epoch (p>0.05). We tested the robustness of these results by detecting that a comparable effect was still present in subpopulations of neurons selected using different criteria (Dn > 5% and Dn > 20%; supplementary Table S1 and corresponding description).

Figure 5 (A and B) displays the pattern of activity from four representative neurons from DLPFC and PMd. Figure 5A shows the temporal evolution of the activity of two different DLPFC neurons during the time around the Pair Onset (upper panels) and the time around the Movement Onset respectively (lower panels). In these neurons, the average neuronal activity during the corresponding epochs was higher for the right position of the target and increased with increasing SDist. Figure 5B displays a similar plot for two PMd neurons. The first of the PMd neurons (upper panels) displays a preference for the left position of the target during the Delay epoch that increased with the growth of SDist, while the second neuron (lower panels) displays a preference for the right position of the target during the time around the movement onset, but this preference was not modulated by the SDist.

**Figure 5:**
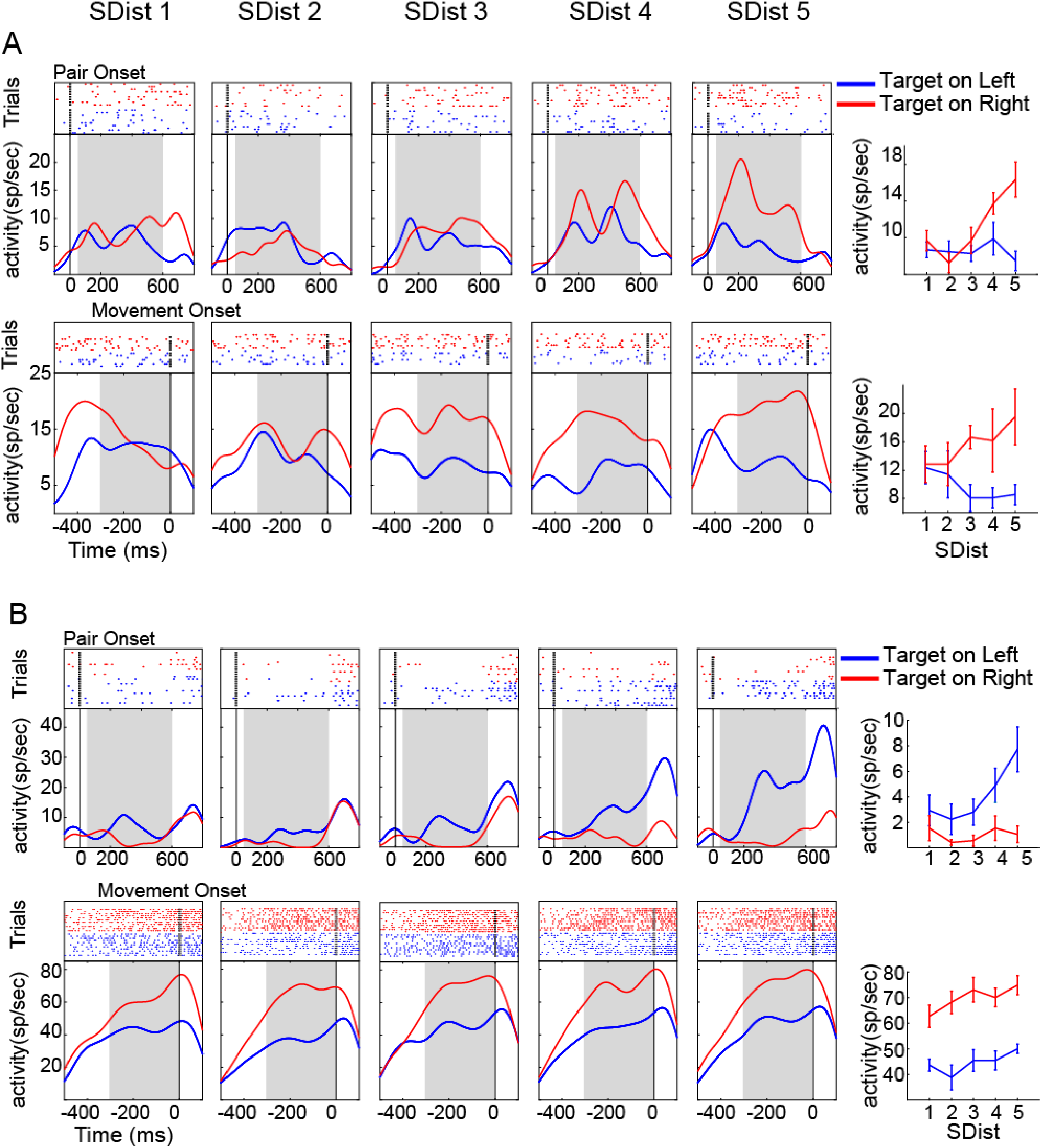
Modulation of activity in DLPFC and PMd neurons during pairs comparisons. Time evolution of the neuronal activity of four different neurons recorded from DLPFC (Panel A) and PMd (Panel B), exhibiting a target position selectivity during the Delay epoch (from 100 ms before to 800 ms after the pair onset; first row) and Reaction Time epochs (from 600 ms before to 200 ms after the movement onset; bottom row), respectively. For comparisons at each SDist, the raster plots (upper part) and the corresponding spike density functions (lower part) are plotted for correct trials grouped according to left (blue) and right (red) target positions. The rightmost panels show the mean spike rate as a function of SDist, calculated during the Delay and RT analysis epochs (shaded area). Both the DLPFC example neurons exhibit a preference for the right target position (red) and an increasing difference in neuronal activity for the left and right target positions with increasing SDist. In contrast, only one of the PMd example neurons (top one) modulated its preference for the left position with the rising SDist.

We also observed that the DLPFC example neuron exhibiting a target position selectivity during the RT epoch (Figure 5A, bottom panel) displays a decreasing spike rate for the left target position at higher SDist (easier target discrimination). We further evaluated whether this effect was shown by more neurons in the DLPFC population using a linear regression analysis, and we observed that 21/141 (14.8%) neurons studied by us in this area exhibited a negative correlation of neuronal activity with the SDist during at least one of the epochs.

We further asked how many neurons contributed to the encoding of the same target position at each level of difficulty and whether the contribution of a single neuron emerged only for specific levels of difficulty. We observed that a small proportion of DLPFC neurons (8% of all the target position selective neurons) exhibited a preference for the same target position at every difficulty level of the task, while in the PMd, we quantified that 41% of all the target-selective neurons maintained the same directional preferences at every SDist.

To summarize, easier pair comparisons in the TI task are related to the recruitment of a higher percentage of target position selective neurons than for more difficult comparisons, both in the DLPFC and PMd. In addition, the spatial selectivity was found to be stronger for easier pair comparisons. However, in PMd neurons, this trend occurred only during the Delay epoch; in the RT epoch, this selectivity was encoded with comparable strength at each level of difficulty.

### The selectivity of the target position occurs earlier in the DLPFC than in the PMd

The previous results indicate that the target position selectivity in both the DLPFC and PMd was influenced by the task difficulty during different epochs of the task. Here, by using an ROC analysis, we explored the time evolution of spatial selectivity of the target position to determine the time at which it occurred in the selected DLPFC and PMd neurons. Figures 6A and 6B show the temporal evolution of the ROC in the time around the pair onset in 47 neurons recorded from the DLPFC and 55 neurons from the PMd of Monkey 1 obtained from the recording sessions with comparable RTs (see methods). Neurons in each plot were sorted according to the time of target position selectivity (see methods for more details). The top histograms on each plot illustrate the distribution of the neuronal estimation of target position selection latencies for each SDist and the corresponding average values. We tested whether the latency in the emergence of the spatial selectivity differed between the two brain areas and whether it depended on the task difficulty by quantifying the probabilities of observing these latencies during the initial 75% (<900 ms) of the distributions at each SDist. We observed a significantly higher probability of finding shorter spatial target selection latencies in the DLPFC than in the PMd (Figure 6C - Kolmogorov Smirnoff test; p<0.05).

**Figure 6.**
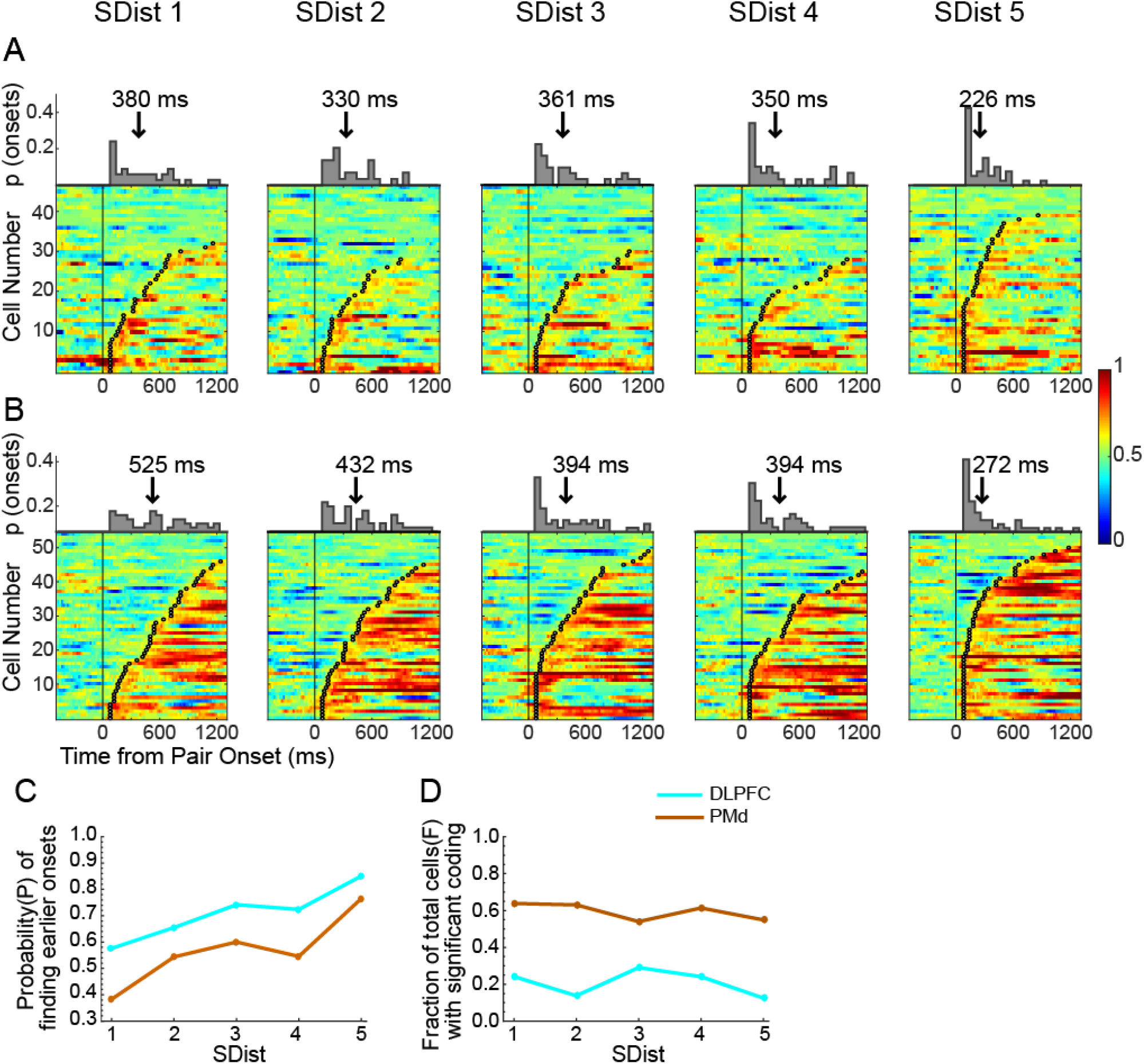
Estimate of the time of target position selectivity in the DLPFC and PMd neuronal population from Monkey 1. **A-B.** Time course of target position selectivity, measured as auROC values between the two target positions, in the time from 500 ms before the pair onset to 1300 ms after it, at each SDist from Population of DLPFC neurons(A) and PMd neurons(B). In each plot, the neurons are aligned sorted to the achievement of the set criterion (crossing the threshold value of 0.6 for 60 consecutive ms), highlighted by the black circles in each row. The histograms on the top show the distribution of these onsets of discrimination between target positions over trial time, and their mean is represented by a vertical arrow for each SDist. **C.** The probability of observing the target selection onsets in the plots (A and B) before 900 ms after the pair onset (75% of the distribution) as a function of SDist in DLPFC and PMd neurons. **D.** Proportion of total DLPFC and PMd cells in Monkey 1 significantly coding the target position for more than 70% of the total time.

Furthermore, we found that the average target position selectivity latency across all conditions was significantly longer in the PMd than in the DLPFC (DLPFC: 329.2 ms; PMd 403.5 ms; t test: t (398) = 2.4079; p<0.05). The probability of observing shorter latencies was also found to be higher with the increase in SDist in both areas. We fitted these trends with a linear model and observed a goodness of fit of 0.78 in DLPC and 0.91 in PMd. Additionally, we detected that the proportion of cells showing sustained encoding for the target position (Figure 6D) was greater in the PMD than in the DLPFC (Kolmogorov Smirnoff test; p<0.01). This effect did not depend on SDist (goodness of fit: DLPFC R^2^ = 0.08; PMd R^2^ =0.53).

These results were confirmed by comparing the evolution of target position selectivity in the neurons recorded from the DLPFC of Monkey 2 (79 neurons) and PMd of Monkey 3 (119 neurons) during the recording sessions with comparable RT (Figure 7A and 7B) across the same trial epoch. We detected a higher probability of finding lower values of target position selection latencies for the DLPFC than for the PMd (figure 7C; Kolmogorov Smirnoff test, p<0.01), with a significant modulation by the SDist in both brain areas (goodness of fit: DLPFC R^2^ = 0.76; PMd R^2^ =0.82). Correspondingly, the mean latencies revealed a significantly earlier target position selectivity in the DLPFC than in the PMd (DLPFC: 338.0 ms; PMd 552.8 ms; t test: t (741) = 9.2980; p<0.05). The fraction of cells encoding the target position was significantly higher in PMd (figures 7D and 7E; Kolmogorov Smirnoff test, p<0.05) than in DLPFC in the current comparison, with no significant modulation by the SDist (goodness of fit: DLPFC R^2^ = 0.02; PMdR^2^ =0.49).

**Figure 7.**
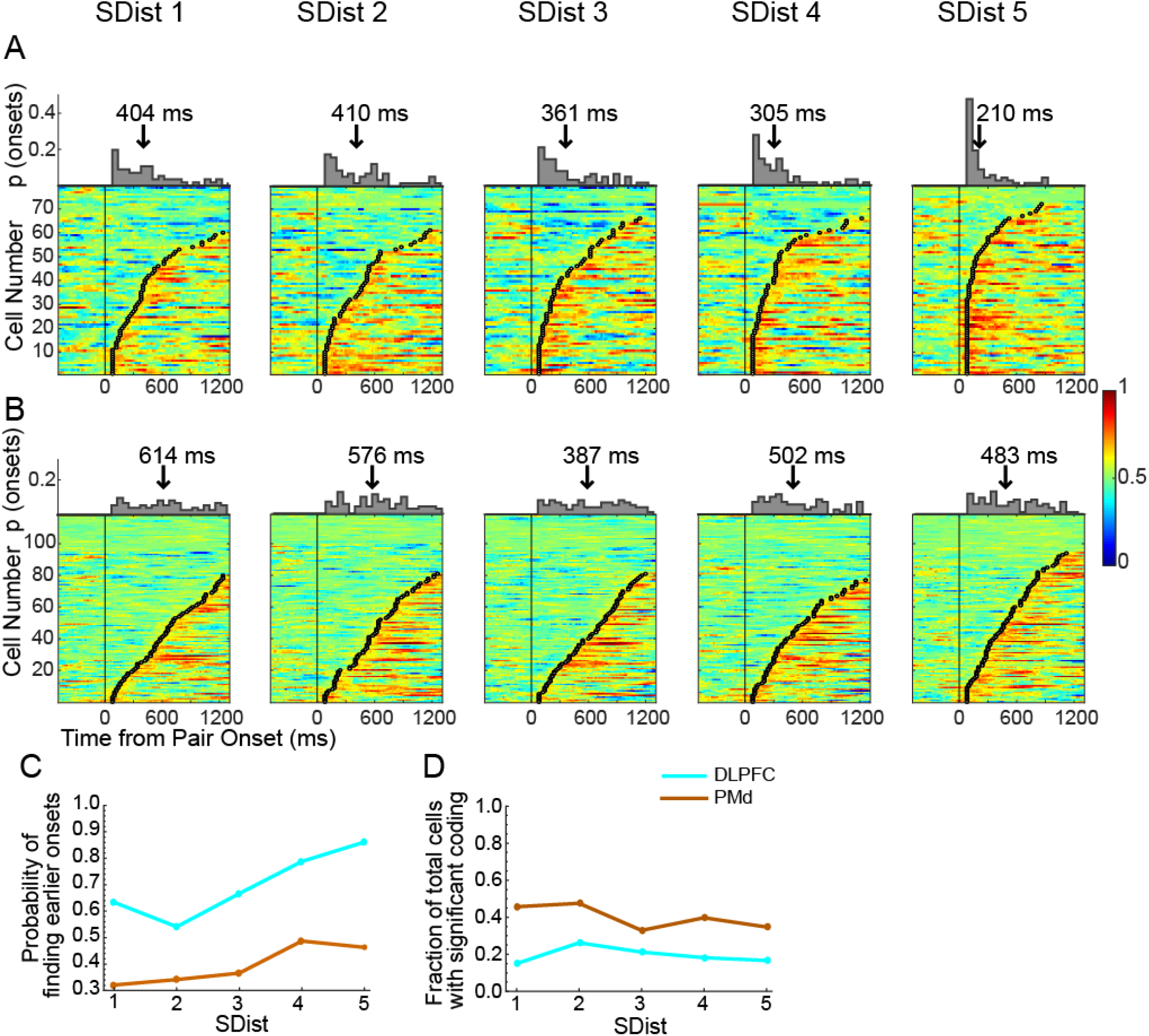
Estimate of the time of target position selectivity in neuronal populations from DLPFC of Monkey2 and PMd of Monkey3. Time course of target position selectivity, measured as auROC values between the two target positions, from the duration of 500ms before the pair onset to 1300 ms after it for each SDist for DLPFC neurons of Monkey2 (A) and PMd neurons of Monkey3 (B). C and D: Comparison of the probabilities of finding shorter target selection onsets and the fraction of cells exhibiting significant coding, respectively.

Collectively, these results indicate that both the DLPFC and PMd contribute to target position discriminability decisions in inferential tasks. In both areas, the latencies and the durations of neuron involvement in spatial selectivity depended upon the degree of task difficulty in detecting the target. More importantly, the two areas partake their role in decision making with earlier involvement in target position coding of DLPFC neurons over PMd neurons. PMd neurons maintained the target position encoding for a longer time, with a higher fraction of cells exhibiting sustained coding.

## Discussion

Building mental models by linking isolated events or situations allows appropriate decision making or conceptualizing new knowledge by relying on linked information (Acuna et al., 2002b; Treichler et al., 2003; Jensen, 2017). The spatial structures of these models have been studied in humans and animals (Constantinescu, 2016; Jensen et al., 2019; Sheahan et al., 2021) to assess how linked information could be mapped. Recently, it has also been demonstrated how spatially organized models allow the generalization of knowledge across contests (Sheahan et al., 2021) and which computational mechanisms subtend these functions (Brunamonti et al., 2016; Mione et al., 2020). The transitive inference task is one of the experimental approaches used to investigate behavioral and neuronal modulations during this form of decision-making in many brain areas, including frontoparietal and hippocampal regions (Acuna et al., 2002a; Zeithamova et al., 2012; Basile et al., 2020). The recent developments of TI tasks in monkey neurophysiological studies have allowed us to investigate the neuronal correlates of this function with improved spatial and temporal resolution (Brunamonti et al., 2016; Mione et al., 2020; Munoz et al., 2020). A view on cortical neuronal processing subtending the manipulation of spatial models of linked information will provide data to validate or refine the biological reliability of the computations modeled by artificial neural networks (Sheahan et al., 2021).

By relying on this approach, we addressed the question of how the difficulty in comparing pairs of items of a learned ranked list modulates the activity of single neurons of both the DLPFC and PMd, two brain areas of the circuit involved in transforming abstract goals in spatially oriented actions (Pezzulo and Cisek, 2016; Grafton and Volz, 2019). The occurrence of the symbolic distance effect in the test session supports our working hypothesis that the symbolic distance effect is consequent to the difficulty arising while making a decision between the rank of two items closely located on the mental representation (Brunamonti et al., 2016; Mione et al., 2020). The analysis of the neuronal data revealed that the degree of difficulty in pair comparisons influenced the spatial selectivity of both DLPFC and PMd neurons. These results confirm our previous observation that the symbolic distance effect modulates the spatial selectivity of PMd neurons (Mione et al., 2020) and document comparable effects in DLPFC neurons.

More specifically, 89% of the DLPFC neurons showed a target position preference, responding with higher levels of activity when the target item was specifically presented at one side of the screen, at least in one of the epochs of the analysis. The proportion of DLPFC neurons preferring the left (45%) and right (54%) directions was comparable and in line with previous studies showing comparable proportions of DLPFC neurons recorded in monkeys performing a motion detection task, while the symbolic distance effect modulated the task difficulty (Lennert and Martinez-Trujillo, 2013). Similar proportions of neurons were observed in PMd (92% neurons selective for the target location; 48% left preference; 52% right preference).

Within this population of target position-selective neurons, we observed an increasing number of neurons exhibiting a preference for the target position in both areas, as the degree of difficulty in distinguishing between the target and the non-target decreased (Figure 3-4). However, the distribution of the target position-selective neurons in the two epochs of analysis, as well as their corresponding level of activation, was modulated differently in the two brain areas. More specifically, in the DLPFC, task difficulty modulated the magnitude of neuronal activity both at the time of target presentation and during the time preceding the onset of movement (Figure 3C), while in the PMd, task difficulty modulated the magnitude of neuronal activity only at the time immediately following pair presentation. For instance, although PMd neurons encoded the target position in the time preceding the movement onset, the magnitude of the encoding was not modulated by the task difficulty. These results reveal that the activity of PMd is more sensitive to the degree of difficulty of the task soon after the pair presentation but not in the later part of the trial and suggest that the neuronal activity of this area was modulated by the uncertainty in target selection only during the earlier part of the trial. In the time immediately preceding movement onset, the neuronal activity of PMd at lower symbolic distances was comparable to that observed at higher symbolic distances, indicating that the target was already selected at this time.

We also observed differences in the DLPFC and PMd neuronal populations in the way they encoded the effect of task difficulty. In both brain areas, we observed an increasing proportion of neurons recruited for easier pair comparisons; however, while in PMd, approximately 40% of neurons were engaged in encoding the target position at different SDists, only 8% of DLPFC neurons kept encoding the same target position at each SDist. These results reveal that the neuronal population encoding the task difficulty was more heterogeneous in the DLPFC than in the PMd, suggesting a higher level of mixed selectivity properties of this area. Additionally, temporal analysis of target position encoding revealed earlier involvement of DLPFC neurons than PMd neurons. Furthermore, PMd cells, after an initial encoding for the target position that depended on task difficulty, kept encoding the target position with higher sustained activity than DLPFC. However, the fraction of cells exhibiting sustained activity was not modulated by task difficulty in either brain area (Figure 6D-7D).

All these results are in line with the hypothesis that PMd is mainly involved in managing the task variables related to motor preparation, while DLPFC activity appears more dependent on abstract variables, such as task difficulty (Yamagata et al., 2012).

However, when performing this last analysis, we needed to consider the different RTs between the three monkeys. This control led us to obtain two separate estimates of the target position encoding time in the DLPFC and PMd, one performed by using the brain activity of the same monkey and the other on the brain activity recorded from different monkeys. Even though in both cases we detected an earlier encoding of target position in DLPFC than in PMd, we observed a larger difference in the time lag in the estimate relaying on the sessions coming from two different monkeys (74.3 ms vs 214.7 ms), likely due to the individual differences. To obtain a reliable estimation of the target selection time of the two brain areas, a simultaneous recording from both the DLPFC and PMd would be preferred. A comparable pattern of neuronal activity in the PFC and PMd evolving with different timings has been observed during the execution of tasks requiring the encoding of abstract rules in matching to sample or quantity comparison tasks (Wallis and Miller, 2003; Vallentin et al., 2012), goal selection in conditional visuomotor tasks (Yamagata et al., 2012) or visual categorization tasks (Cromer et al., 2011). In all these tasks, the PMd cortex was observed to use the task variables to form a motor plan during the time of movement onset and poorly engaged in the encoding task variable not linked to a motor decision (Cromer et al., 2011; Vallentin et al., 2012; Yamagata et al., 2012).

In contrast, the DLPFC is a higher-order brain area involved in encoding abstract variables of decision, such as decision rules (Kim and Shadlen, 1999; Bongard and Nieder, 2010), strategy (Genovesio et al., 2006), behavioral goals (Yamagata et al., 2012; Falcone et al., 2016) or the rank order representation of related information (Brunamonti et al., 2016). There is agreement that the DLPFC sends top-down information to downstream motor areas, such as the PMd, to convert the processing of these variables into motor decision commands (Barbas and Pandya, 1987, 1989; Luppino et al., 2003). According to this model, the involvement of the DLPFC in decision making should occur earlier than the PMd.

However, an earlier involvement of the PMd compared to the DLPFC in encoding the characteristics of the task has also been found (Wallis and Miller, 2003; Cromer et al., 2011). These authors ascribe this early involvement to the familiarity of the monkeys with the task demands and stimuli, which would allow solving the task without the need for an abstract manipulation of the information, a competence supported by PFC (Muhammad et al., 2006). Our experimental protocol hinders such an eventuality. For instance, the monkeys were required to learn the ranked series from a set of never-experienced items randomly selected at the beginning of each experimental session. Thus, this protocol aimed to prevent familiarity with the task stimuli and increase the involvement of the PFC in driving the decision-making-related activity of PMd. In contrast to the discussed results (Wallis and Miller, 2003; Muhammad et al., 2006; Cromer et al., 2011), the time course of the decision-making-related activity of the PFC and PMd observed here fits more with the proposed anatomo-functional model of the relationship between the two brain areas (Barbas and Pandya, 1987, 1989; Luppino et al., 2003).

A further possible role played by the PFC is the orientation of spatial attention (Lennert and Martinez-Trujillo, 2011; Messinger et al., 2021) before the formation of the proper motor plan. Symbolic distance has been observed to modulate the time and magnitude of target selection-related neuronal activity by decreasing the intensity of the neuronal response when the distracting item is easily distinguishable from the target (Lennert and Martinez-Trujillo, 2011). In our data, we observed such modulation of neuronal activity in the DLPFC only in a minority proportion of neurons (14.8%), suggesting that attentional allocation would also act with a different mechanism.

Our results are in accordance with several lines of research on perceptual decision-making in monkeys, converging on the hypothesis that the resolution of the ambiguity of perceptual stimuli supports the selection between competitive motor actions simultaneously available (Cisek and Kalaska, 2005, 2010; Klaes et al., 2011; Kubanek et al., 2013; Shushruth et al., 2018). Enacting this process, higher-order association areas continuously support the motor system on this decisional computation (Pezzulo and Cisek, 2016; Shushruth et al., 2018). Here, we show that this mechanism can be valid even when ambiguity relates to the representation of information stored in memory.

In summary, the present results support the hypothesis of a hierarchal organization between brain areas in which the DLPFC encodes the variables for decision processes and PMd uses this information to transform abstract decisions in motor programs.

## Notes

### Competing Interest Statement

The authors have declared no competing interest.

### Summary of Updates

A piece of more detailed information on the Methods has been provided in this revised version of the manuscript, along with the supplemental information to support some of the results.

