## Supplementary Information for "Different Contribution of the Monkey Prefrontal and Premotor Dorsal Cortex in Decision Making during a Transitive Inference task"

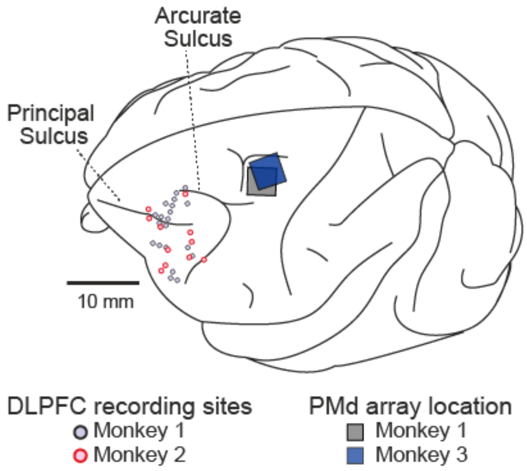


**Figure S1**: **Recording locations**

Recording sites and array location during DLPFC (Monkey 1 and Monkey 2) and PMd (Monkey 1 and Monkey 3) experiments.

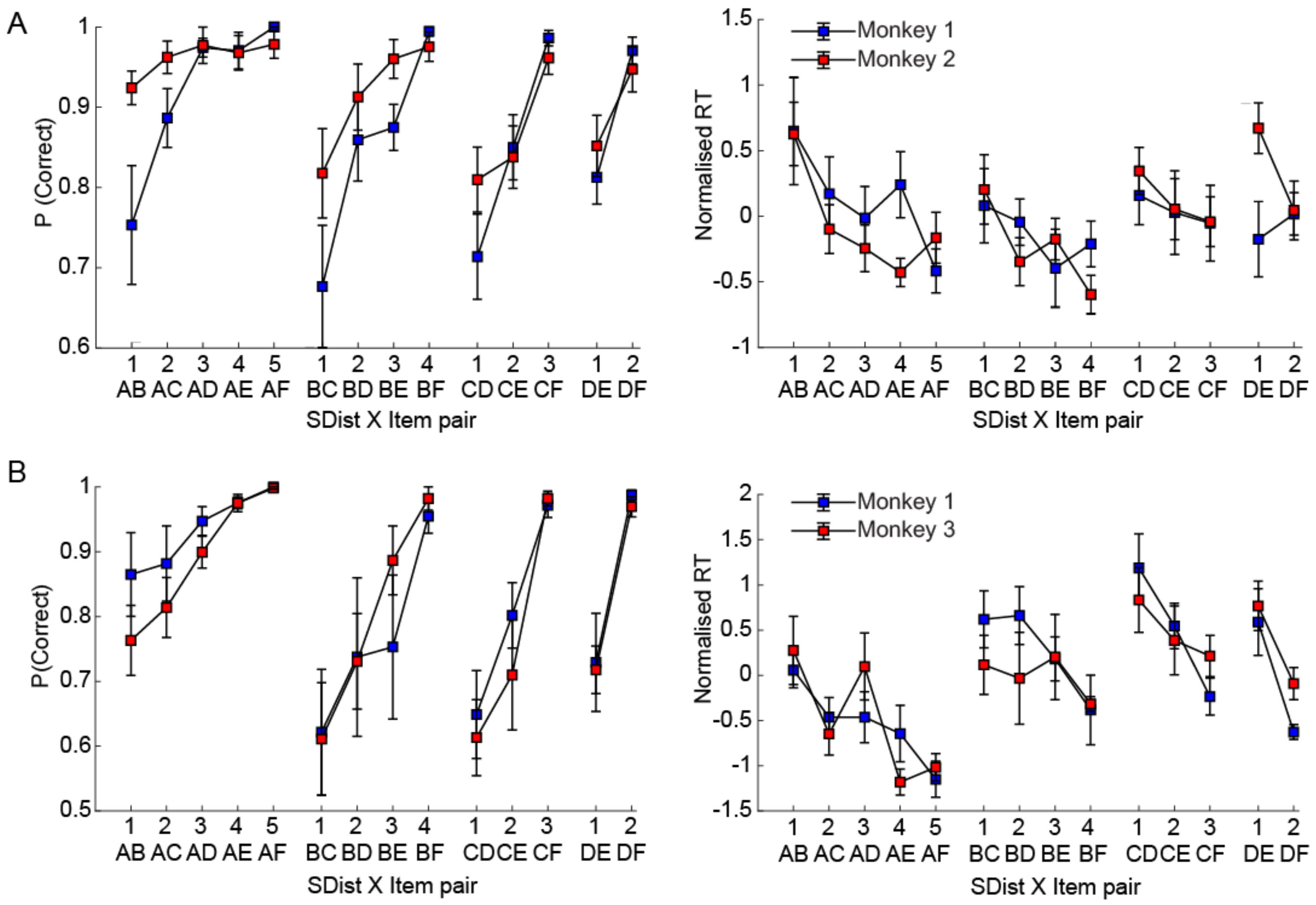

**Figure S2: Performance and normalized reaction times for each paired comparison at different symbolic distances.**

The performance and RTs in selecting the correct target during **A)** DLPFC (Monkey 1: n=14; Monkey 2: n=12) and **B)** PMd (Monkey 1: n=7; Monkey 3: n=7) sessions are represented for each pair organized according to the symbolic distance characterizing each pair. The vertical bars represent the standard error of mean across the sessions. The proportion of correct choices and the normalized RT for each target item was significantly correlated by the symbolic distance from the presented non-target item (linear regression analysis, all p<0.05), except for the RTs of Monkey 1 DLPFC recordings in comparison of item D; and RTs for comparisons of items B and C in the PMd recordings from Monkey 3 (p>0.05). The emergence of symbolic distance effect on the performance and RTs for individual items, especially in the case of the item A, which was always rewarded, supports the hypothesis of a mental representation of the ranked items.

| Mean abs ($D_n$) | SDist1 | SDist2 | SDist3 | SDist4 | SDist5 | Linear Regression Coefficients and statistics |
|---|---|---|---|---|---|---|
| **DLPFC** | | | | | | |
| Delay Epoch (5%) | 0.78 | 0.90 | 1.08 | 1.16 | 1.19 | $D_n$=0.10(SDist)+0.70;p<0.001 |
| RT Epoch (5%) | 0.87 | 0.89 | 1.08 | 1.15 | 1.04 | $D_n$=0.05(SDist)+0.83; p<0.001 |
| Delay Epoch (20%) | 1.36 | 1.36 | 1.52 | 1.56 | 1.54 | $D_n$=0.05(SDist)+1.13; p<0.01 |
| RT Epoch (20%) | 1.39 | 1.43 | 1.55 | 1.66 | 1.68 | $D_n$=0.08(SDist)+1.30; p<0.001 |
| **PMd** | | | | | | |
| Delay Epoch (5%) | 0.84 | 0.96 | 1.00 | 1.01 | 1.31 | $D_n$=0.09(SDist)+0.73; p<0.001 |
| RT Epoch (5%) | 1.08 | 1.01 | 1.09 | 1.10 | 1.19 | $D_n$=0.02(SDist)+1.05; p=0.08 |
| Delay Epoch (20%) | 1.13 | 1.47 | 1.57 | 1.52 | 1.63 | $D_n$=0.05(SDist)+1.37; p<0.001 |
| RT Epoch (20%) | 1.40 | 1.36 | 1.39 | 1.43 | 1.46 | $D_n$=0.01(SDist)+1.35; p=0.06 |

**Table S1: Spatial selectivity (mean abs $D_n$) for DLPFC and PMd neurons at different selection thresholds and their correlation with SDist.**

The evaluation of effect of task difficulty on target position selectivity with different selection criteria of ($D_n$>5%) and ($D_n$>20%) revealed higher (DLPFC: 97.1% and PMd:97.2%) and lower (DLPFC: 64% and PMd: 84%) percentages of target selective neurons respectively, when compared to the target position selective population obtained using a criterion of ($D_n$>10%; DLPFC: 89.3% and PMd: 91.9%). Table S1 reports the mean absolute values of $D_n$, showing the strength of overall target position selectivity at each SDist. A linear regression analysis revealed that the strength of target position selectivity is significantly correlated with the task difficulty during both the analysis epochs in DLPFC neurons but only during the delay epoch in the PMd neurons even with these selection criteria. Overall these results confirm that at a population level SDist influences the spatial selectivity and that this effect can be detected even applying different criterions of selection of populations of neurons.
